# Pre-existing Subclones Determine Radioresistance in Rectal Cancer Organoids

**DOI:** 10.1101/2023.11.02.565315

**Authors:** D. Andel, B.J. Viergever, N.A. Peters, D.A.E. Raats, S.J. van Schelven, M.P.W. Intven, M. Zandvliet, J. Hagendoorn, I.H.M. Borel Rinkes, O. Kranenburg

**Affiliations:** Department of Surgical Oncology, University Medical Center Utrecht, Cancer Center, Utrecht, The Netherlands; Laboratory for Translational Oncology, University Medical Center Utrecht, Cancer Center, Utrecht, The Netherlands; Department of Radiation Oncology, University Medical Center Utrecht, Cancer Center, Utrecht, The Netherlands; Department of Clinical Sciences - Companion Animals, Faculty of Veterinary Medicine, Utrecht University, Utrecht, the Netherlands; Utrecht Platform for Organoid Technology, Utrecht University, Utrecht, The Netherlands

**Keywords:** Single-cell DNA sequencing, tumor evolution, patient-derived cancer organoids, intratumor heterogeneity, chromosomal instability, radiation therapy resistance

## Abstract

More than half of all cancer patients receive radiation therapy, but resistance is commonly observed. Currently, it is unknown whether resistance to radiation therapy is acquired or inherently present. Here, we employed organoids derived from rectal cancer and single-cell whole genome sequencing to investigate the long-term evolution of subclones in response to radiation. Comparing single-cell whole genome karyotypes between unirradiated and irradiated organoids revealed three patterns of subclonal evolution: (i) subclonal persistence, (ii) subclonal extinction, and (iii) subclonal expansion. Only organoids in which subclonal shifts occurred (i.e., expansion or extinction) became more resistant to radiation. Although radioresistant subclones did not share recurrent copy number alterations that could explain their radioresistance, resistance was associated with reduced chromosomal instability; an association that was also observed in 529 human cancer cell lines. These data suggest resistance to radiation is inherently present and associated with reduced chromosomal instability.

## Introduction

Radiation therapy constitutes a cornerstone of cancer treatment, but resistance poses a major clinical challenge. For example, more than 70% of patients with esophageal (1) and rectal cancer (2) receiving chemoradiation therapy have residual disease. Gaining a better understanding of the mechanisms underlying resistance could potentially optimize the efficacy of radiation therapy, and aid in the identification of patients who are most likely to respond.

Genomic studies in various malignancies have revealed that resistance to systemic therapy can arise through *de novo* (e.g., treatment-induced) genomic aberrations (3–5), or may be inherently present, such as through the selection of (rare) pre-existing subclones (3,6,7). Deciphering these mechanisms of resistance is important because if it is indeed inherently present, response to therapy could potentially be predicted by analyzing pretreatment biopsies.

Many of these studies have used bulk genome sequencing to infer subclonal relationships in response to cancer treatment. However, bulk genomic sequencing lacks the ability to resolve intratumor heterogeneity at a high resolution. As such, the existence of pre-existing rare subclones may go unnoticed. In contrast, single-cell whole genome sequencing has emerged as a powerful tool enabling reconstruction of phylogenetic trees (8) and detection of rare subclones (9,10).

In the field of radiotherapy, research on this topic has been impeded by challenges in obtaining serial patient samples over time and the lack of suitable *in vitro* models.

Recently, organoids have emerged as robust cancer models that can accurately predict clinical responses to radiation therapy in rectal cancer (11,12). Additionally, organoids recapitulate the genomic intratumor heterogeneity at the single-cell level (13), making them a powerful model for studying resistance to radiation.

Currently, it remains unknown if resistance to radiation therapy is treatment-induced through the generation of *de novo* genomic aberrations or inherently present via the selection of (rare) pre-existing subclones. To address this question, we utilized patient-derived organoids from rectal cancer and employed single-cell whole genome sequencing to track subclonal evolution in response to radiation.

## Results

### Patient-derived organoids from rectal cancer display heterogeneous responses to radiation therapy

Eight patient-derived organoids from primary rectal cancers were established, covering all non-in situ AJCC stages. Table S1 provides detailed information on the basic clinical and tumor characteristics of these organoids. Out of eight organoids, two were derived from patients who received neoadjuvant chemoradiotherapy, while the remaining six were from treatment-naïve patients. The median overall survival from the time of surgery was 1,857 days, and the median progression-free survival was not reached. Next, a high-throughput radiation response assay was utilized, which closely mimics radiation response parameters of the classic clonogenic survival assay in cell lines (14,15), and was adapted for compatibility with 3D organoid models (Methods). Briefly, organoids were grown for three days in BME microdroplets and exposed to a range of clinically relevant radiation doses. After seven days, post-radiation cell viability was assessed using CellTiter-Glo 3D, which quantifies intracellular ATP. Differences in sensitivities to radiation therapy were observed between models derived from different patients (Fig. 1a). Based on the median values of the area under the curve, three organoids were found to be consistently resistant (HUB005, HUB183 and HUB015) or sensitive (HUB106, HUB197 and HUB062) to radiation in multiple independently performed experiments (Fig. 1b).

**Figure 1.**
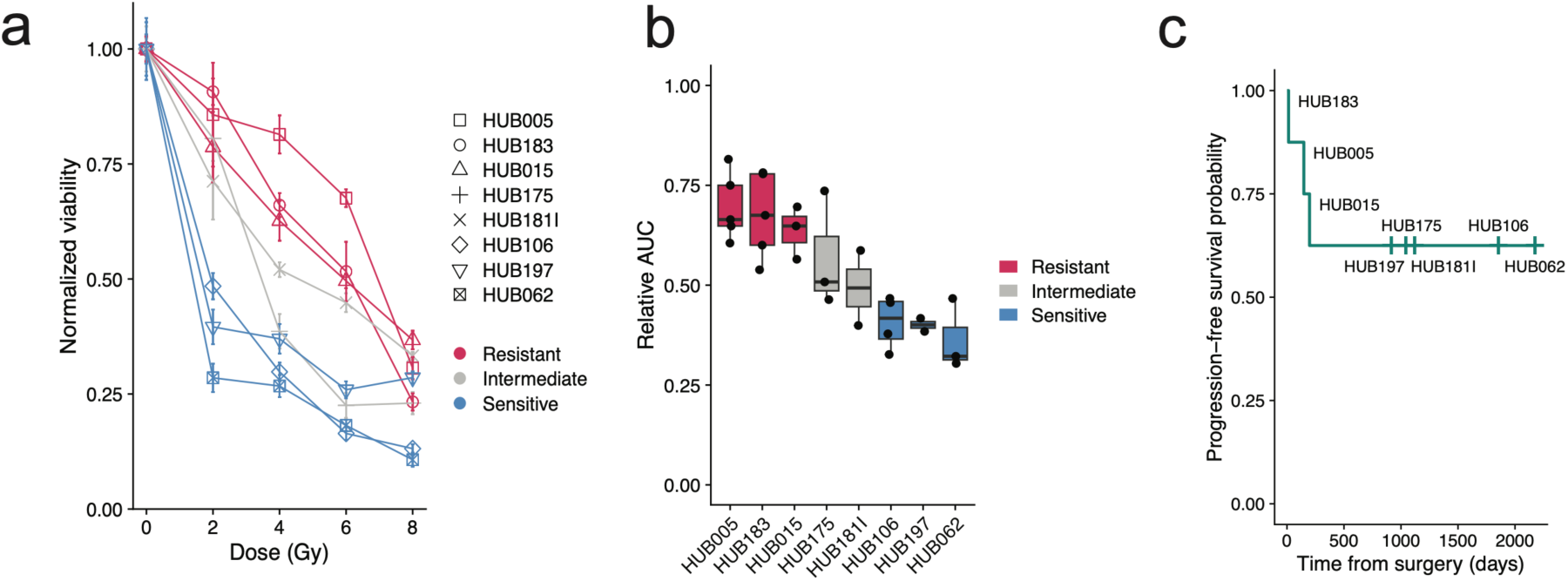
Heterogeneous *in vitro* response to radiation associates with progression-free survival a, Representative dose response plot showing responses (normalized to 0 Gy) of eight rectal cancer-derived organoids. **b,** Combined analysis of the relative area under the curve (AUC) of multiple independent experiments (each dot) reveals low inter-experiment variability and identifies three radioresistant organoids (red) and three radiosensitive organoids (blue). **c,** Progression-free survival from the time of surgery (and organoid harvest) for each rectal cancer organoid demonstrates patients corresponding to resistant organoids progressed early, while patients from whom sensitive organoids were derived had no progression (censored, vertical line).

*In vitro* organoid chemoradiation responses correlate with clinical progression-free survival (11,16). In this study, a statistical correlation between *in vitro* radiation resistance and progression-free survival could not be performed due to censoring of data (i.e., no progression occurred). However, patients with *in vitro* radiation resistant organoids (HUB005, HUB183, HUB015) all progressed early after surgery (after 148, 14 and 198 days, respectively), while patients corresponding to radiation sensitive organoids (HUB106, HUB197, HUB062) had no progression after 1857, 917 and 2175 days, respectively (Fig. 1c). Based on these data, three radioresistant organoids (HUB005, HUB183, and HUB015) and three radiosensitive organoids (HUB106, HUB197, HUB062) were selected for further analysis.

### Radioresistant and radiosensitive organoids have colorectal cancer-specific driver mutations and copy number profiles

To identify mutations in driver genes common to colorectal cancer, targeted or whole exome sequencing was performed. This revealed mutations in *TP53* (6/6), *APC* (4/6), *SMAD4* (1/6), *PTEN* (1/6) and *PIK3CA* (1/6) (Supplementary Tables 2 and 3). Sparse (∼0.1x) single-cell whole genome sequencing showed that most of the organoids (5 out of 6) exhibited high frequencies of arm or chromosome-level copy number alterations, which is also frequently observed in samples from colorectal patients (Fig. 2a, e, Fig. 3a, Fig. S1a, e) (17). Analysis of the single-cell karyotype data showed the presence of several colorectal cancer-specific arm-level copy number gains, such as 1q, 7, 8, 13 and 20 p and q as well as deletions in 18q and 8p, consistent with patient data from the TCGA database (17). HUB015 was the only organoid that was largely diploid, which is apparent in approximately 16% of colorectal cancers (Fig. 3e) (17,18). HUB197 displayed cells that likely arose from a whole-genome duplication event, as these cells had exactly twice the ploidy of another set of cells within the organoid culture (Fig. 3a) (19,20).

**Figure 2.**
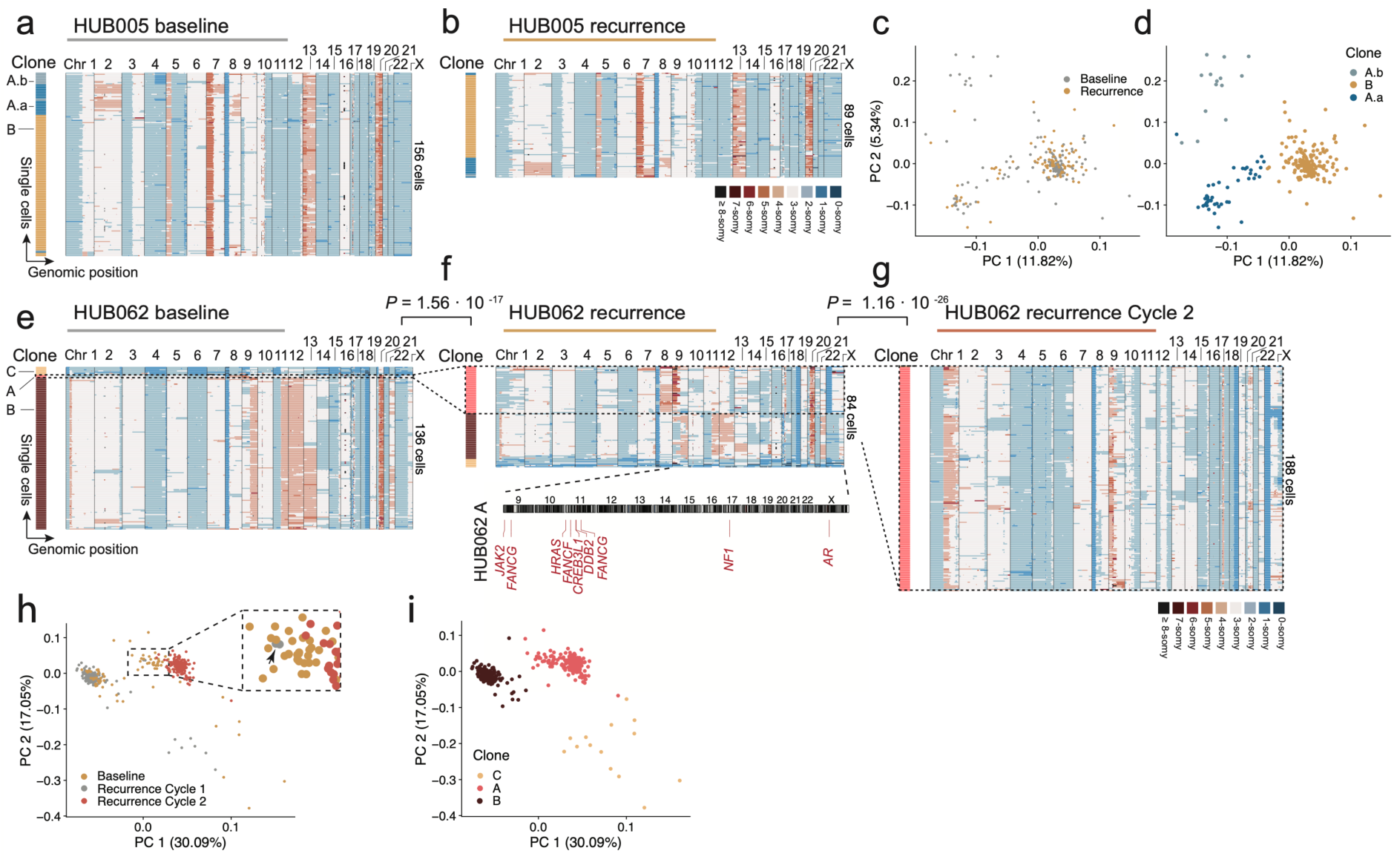
Subclonal evolution in response to irradiation in persister and expansion organoid. a, b, Clustered heatmaps of single cell copy number profiles of HUB005 (resistant, persister) at baseline (**a**) and after recurrence following 10 Gy of irradiation (**b**). **c, d,** PCA plots of single cell copy number profiles from the baseline and recurrence populations. Colors indicate treatment status (**c**) or subclone (**d**) as defined using k-means clustering. **e, f, g**, Cluster heatmaps of single cell copy number profiles of HUB062 (sensitive, expander) at baseline (**e**), after recurrence following 10 Gy (**f**) and after recurrence following another cycle of 10 Gy (**g**). **f,** Lower panel indicates the karyotype plot of expansion clone HUB062A with amplified oncogenes specific to HUB062A (when compared to HUB062B and HUB062C). **h, i,** PCA plots of single cell copy number profiles from the baseline, recurrence and recurrence cycle 2 populations. Inset in **h** localizes two cells of subclone HUB062A at baseline (arrow head). Formal subclones (Methods) are depicted with capital letters, while minor subpopulations within clones are indicated with a suffix (.a, .b etc.). *P*-values are from Fisher’s exact test.

**Figure 3.**
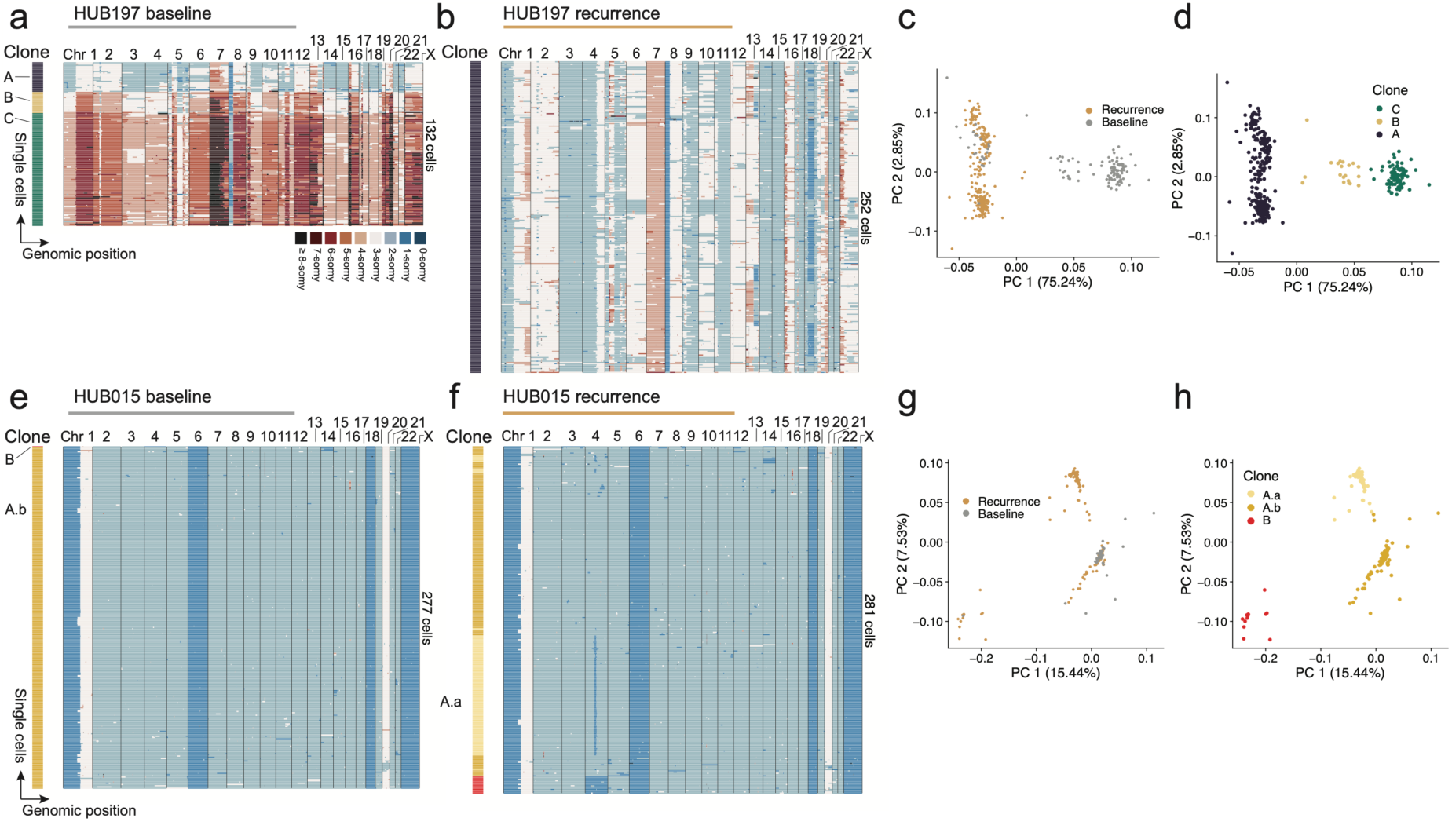
Subclonal evolution in extinction and persister organoid. a, b, Single cell copy number heatmaps of HUB197 (sensitive, extinction organoid) at baseline (**a**) and after recurrence following 10 Gy of irradiation (**b**). **c, d,** PCA plots of single cell copy number profiles from the baseline and recurrence populations with colors indicating treatment status (**c**) or subclone (**d**). **e, f,** Cluster heatmaps of single cell copy number profiles of HUB015 (resistant, persister) at baseline (**e**) and after recurrence following 10 Gy (**f**). **g, h,** PCA plots of single cell copy number profiles from the baseline and recurrence populations. Formal subclones (Methods) are depicted with capital letters, while minor subpopulations within clones are indicated with a suffix (.a, .b etc.).

### Distinct modes of subclonal dynamics in response to radiation

Cancer cells may survive therapeutic pressures through inherent or acquired (e.g., treatment-induced) genetic alterations (3–5,7,21,22). To delineate the subclonal evolution in response to radiation, organoids were subjected to either 0 or 10 Gy of radiation. Single-cell whole genome karyotypes of unirradiated organoids (referred to as ‘baseline’) were compared to irradiated organoids after they were amenable to passaging (referred to as ‘recurrence’). Recurrence following 10 Gy was observed after 31 ± 12 days in resistant organoids and after 49 ± 24 days in sensitive organoids. Next, the genetic information of single cells obtained at baseline and at recurrence was plotted on the same graph using principal component analysis. This way, cells with similar copy number alterations before and after treatment are part of the same subclone and thus cluster together. This allowed detection of pre-existing subclones that persisted or expanded upon treatment. Conversely, posttherapy cells with treatment-induced *de novo* genomic aberrations will form distinct clusters that are not in the vicinity of pretherapy cells. It was reasoned that cells with a set of similar copy number alterations should only be considered as part of a ‘formal’ subclone when the copy number alterations explained more than 10% of the variance within a principal component (Methods).

Using this approach, an average of 2.3 subclones per organoid were identified, and three different patterns of subclonal evolution in response to radiation were observed:

i. subclonal persistence, where the subclones persisted after treatment (Fig. 2a-d, Fig.

3e-h, and Fig. S1a-d), (ii) subclonal extinction, wherein pre-existing subclones died out following treatment (Fig. 3a-d), and (iii) subclonal expansion, wherein a pre-existing subclone expanded (Fig. 2e-i). Evidence for the creation of new subclones after radiation was not found. In fact, although radiation increased the total number of copy number alterations (Fig. S2) (23), *de novo* copy number alterations shared by more than 50% of cells were rare: only in HUB015 a distinctive new deletion spanning 4q21.21 – 4q22.2 was detected (Fig. 3e, f).

Notably, subclonal persistence was observed in resistant organoids (3 out of 3 resistant organoids), while subclonal extinction and subclonal expansion were only seen in sensitive organoids (2 out of 3 sensitive organoids).

For example, HUB005, a resistant organoid, had two subclones (A and B) at baseline; with A subdivided in two minor subclones A.a (4q diploid) and A.b (4q deletion) (Fig. 2a-d). Upon radiation, all subclones persisted in equal proportions (Fisher’s exact test, *P* = 1, Bonferroni corrected). Persistence was also seen in HUB015 (Fig. 3e-h), HUB183 (Fig. S1a-d) and HUB106 (Fig. S1e-h). On the other hand, HUB197 (sensitive), which initially had three subclones at baseline, experienced complete extinction of its two subclones that had whole-genome duplication (*P* = 1.99 · 10^-71^, Fig. 3a-d). Lastly, in HUB062, a radiosensitive organoid, there was a subclone (subclone HUB062A) that constituted only 1.5% of all cells at baseline, but expanded upon radiation to make up 47% of all cells at recurrence (*P* = 1.56 · 10^-17^, Fig. 2e, f). An independent repeat experiment also showed an aggressive expansion of HUB062A, with complete extinction of subclones HUB062B and HUB062C (Fig. S3).

### Subclonal shifts alter sensitivity to radiation

It was hypothesized that organoids that survived therapy and exhibited evidence of subclonal persistence would retain their inherent radioresistance. On the other hand, organoids that showed evidence of subclonal extinction or subclonal expansion were expected to exhibit increased resistance due to shifts favoring radioresistant subclones. Indeed, HUB197 (extinction) and HUB062 (expander) showed increased resistance to radiation compared to their parental counterparts (Fig. 4a). In contrast, the resistance of three persister controls (HUB005, HUB183, HUB106) did not change (Fig. 4b). Moreover, treating the recovered HUB062 again with 10 Gy resulted in complete dominance of the expansion subclone HUB062A in two independent experiments (from 47% to 100%, Fisher’s exact test, *P* < 1.16 · 10^-26^, Fig. 2f, g).

### Copy number alterations associated with radiation resistance

Copy number alterations may harbor oncogenes that confer a survival benefit to cancer cells when amplified or deleted. Resistance-associated copy number alterations were identified by computing consensus copy number alterations for each subclone and selecting only those alterations that were not frequently present in sensitive subclones (Methods). Determining such resistance-specific copy number alterations revealed that expansion subclone HUB062A, when compared to HUB062B and HUB062C had unique copy number amplifications containing oncogenes previously linked to radioresistance, including chromosome-arm 9p (containing proto-oncogenes *JAK2*, *FANCG*), 11p (*FANCF*, *DDB2*) and Xp11.22 - Xq23 (*AR*) (Fig. 2f, lower panel, Table S4). Fanconi anemia complementation group (FANC) family of genes are involved in DNA interstrand crosslink repair pathways (24). The acquired deletion in HUB015 on 4q21.21 – 4q22.2 contained tumor suppressor gene *PTPN13*, coding a protein tyrosine phosphatase, whose inactivation increases invasiveness of cancer cells (25). However, no specific copy number alterations shared by all radioresistant subclones were found, indicating additional mechanisms played a role in how these subclones persisted or expanded.

### Radioresistant subclones have copy number patterns associated with decreased chromosomal instability

To better understand why some subclones showed resistance to radiation, the single-cell karyotype data were carefully reexamined. Radiosensitive subclones within HUB062, HUB106 and HUB197 had more large-to-whole chromosome copy number changes compared to other subclones. Additionally, subclones HUB197B and HUB197C - which did not survive after radiation therapy- exhibited whole-genome duplication (19,20). Both these observations hint towards increased ongoing mitotic chromosome segregation errors within these sensitive subclones. To test this hypothesis, cell-to-cell copy number variability, a measure of chromosomal instability, was assessed within each subclone, as well as arm to whole-chromosome-level copy number alterations. Arm to whole-chromosome-level copy number alterations are suggestive of chromosomal instability due to mitotic errors (26). This analysis revealed that sensitive subclones were indeed more heterogeneous (student’s *t* test, *P* = 1.6 · 10^-2^, Fig. 4c). Moreover, cells within sensitive subclones harbored more copy number alterations, especially at the arm to whole-chromosome-level (Mann-Whitney *U* test, *P* = 1.4 · 10^-63^ and 4.2 · 10^-83^, Fig. 4d).

**Figure 4.**
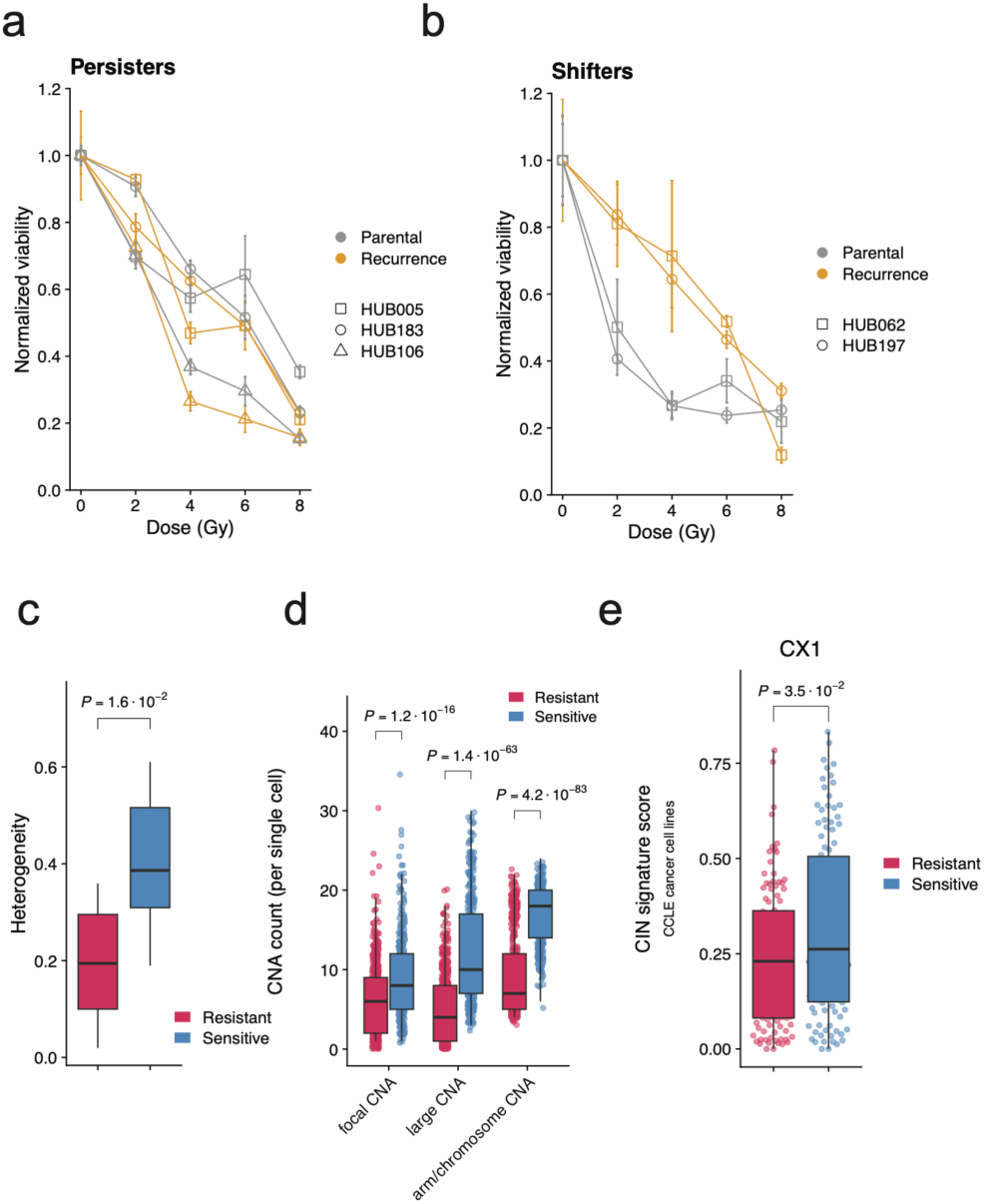
Radioresistance is associated with copy number patterns suggestive of mitotic chromosomal instability a, Representative dose response plot showing responses to irradiation between parental and recurrence lines in organoids where subclonal shifts were apparent (HUB062, HUB197). **b,** Dose response plot in subclonal persisters (HUB106, HUB183 and HUB005). Viability was normalized to 0 Gy. **c,** Heterogeneity score for resistant (red) versus sensitive (blue) subclones. **d,** Copy number alteration counts (stratified into focal, large and arm to whole- chromosome level size) in single cells from resistant or sensitive subclones. **e,** Chromosomal instability signature ‘CX1’ scores from cell lines in the top 20% radioresistant (red) versus the top 20% radiosensitive (blue) cancer cell lines from the CCLE database. CNA: copy number alteration, CIN: chromosomal instability.

Next, recently published chromosomal instability signatures (26) were computed of 529 cancer cell lines from the Cancer Cell Line Encyclopedia (CCLE) of which both radiation sensitivity and copy number data was available (15,27). After correcting for cancer (sub)type, we found that two (2/17) signatures were significantly different when comparing the top 20% radioresistant to the top 20% sensitive cancer cell lines. Signature CX1 – related to arm or whole-chromosome changes due to mitotic errors – was increased in sensitive cell lines (*P* = 3.5 · 10^-2^, Fig. 4e, Table S5), while CX7 unkown aetiology, *P* = 6.6 · 10^-3^) was decreased (26).

## Discussion

In the present study, we tracked subclonal evolution in response to radiation therapy using rectal cancer patient-derived organoids and single-cell whole genome sequencing. We observed either subclonal persistence, subclonal extinction, or subclonal expansion upon radiation therapy, but the creation of new, commonly shared genomic aberrations was rare. Radiosensitive subclones exhibited copy number patterns indicative of mitotic segregation errors. This suggests that subclones may be selected based on decreased chromosomal instability.

The results presented here align with previous studies in patients with breast cancers who are treated with a combination of mitotic inhibitors and anthracyclines, which induce DNA damage. These studies have shown that pre-existing subclones can either expand or go extinct, but new subclones are not generated (6,21). In our study, five out of six analyzed rectal cancer organoids exhibited widespread arm or chromosome-level copy number alterations at baseline (i.e., before irradiation), suggesting chromosomal instability (17). We surmise that chromosomal instability allows the generation of subclones with copy number alterations that confer a survival benefit upon external pressures (28). However, we found no copy number alterations harboring amplified or deleted genes that could comprehensively explain the observed radioresistance of subclones, warranting similar analyses in larger studies in the future.

Although the existence of pre-existing subclones could potentially lead to resistance through their selection and expansion, the data presented here also suggests that increased chromosomal instability is associated with increased sensitivity to radiation therapy. This is supported by studies showing that suppressing chromosomal instability can increase resistance to radiation (29), and that patients with high pretreatment chromosomal instability have better response rates to chemoradiation therapy (30). This association may be explained by the fact that radiation itself induces mitotic segregation errors (29,31), as such overwhelming the cell’s capacity to cope with the resulting genetic abnormalities (32). Interestingly, resistant organoids all showed subclonal persistence, while subclonal shifting (expansion or extinction) was only observed in sensitive organoids. Although radiation can cause numerous copy number alterations through poorly repaired DNA breaks (23), we only observed one case of a newly acquired copy number alteration shared by more than 50% of the cells within a subclone. It appears that radiation-induced alterations are difficult to select for, possibly because other alterations can be harmful to the cell’s survival.

While this study suggests resistance to radiation appears to be primarily determined by genetic factors that are inherently present, resistance on the epigenetic or transcriptional level may be acquired, and these processes do not necessarily exclude each other (6). Cell autonomous factors appear to be important drivers of radioresistance, as indicated by the strong correlation between the radiation response of organoids *in vitro* and clinical response (11,12). Nevertheless, further research on patient samples is necessary to account for the effects of the microenvironment on subclonal evolution.

The findings presented here appear to indicate that resistance to radiation therapy is largely determined by pre-existing radioresistant subclones that either persist or expand, rather than being newly created. This implies that in theory, radioresistance could be predicted by careful analysis of pretreatment biopsies, ideally using techniques that can resolve intratumour heterogeneity at great resolution. Analysis of copy number patterns also indicates that chromosomal instability may be a biomarker for response to radiation therapy.

## Acknowledgements

The authors would like to express their gratitude to Geert Kops and Nico Lansu for their valuable scientific insights and code sharing, and Joshua Peterson and Floris Leendert for their support during the single-cell experiments. Special gratitude goes towards André Wopereis, Wilfred de Vries, Ingrid Boots, Marjolijn Gross, and Masha de Koning-Hoogeboom for their practical insights and assistance with the irradiation of organoids. This research was funded by a private fund.

## Author contributions

D.A. conceived and performed experiments, wrote the code and analyzed the data and wrote the original manuscript. B.J., N.A.P., D.A.E.R and S.J.vS performed experiments. M.P.W.I and M.Z. provided resources. J.H. conceived and supervised. I.H.M.B.R conceived and supervised, and secured funding. O.K. conceived, supervised, wrote the original manuscript, and secured funding. All authors provided expertise and feedback throughout the project and reviewed and edited the manuscript.

## STAR Methods

### Patient-derived tissue and clinical data

Written informed consent was obtained from all patients for the research use of their tissue and processing of clinical data, following the HUB biobank protocol HUB-cancer TcBio#12-09. Clinical data, including pTNM stage, (neo)adjuvant therapies, progression-free survival and overall survival from the time of surgical resection, were extracted by an independent data manager using a custom query. All clinical data accessible to the researchers were fully anonymized. The study was approved by the ethical review board of the University Medical Center Utrecht.

### Patient-derived tumor organoid culturing

Rectal cancer patient-derived organoids were derived and cultured according to previously published protocols (33,34). In brief, organoids were cultured in colorectal cancer culture medium consisting of advanced DMEM/F12 medium (Invitrogen) with supplements as detailed in Supplementary Table 6. Tumor organoids were passaged by dissociating them with TrypLE (Gibco) for 5 minutes, followed by embedding them in a mixture of Basement Membrane Extract (BME; Amsbio) and culture medium in a 3:1 ratio. The resulting tumor organoids were then replated in drops of approximately 10 μl each, in a pre-warmed 6-well plate. To prevent anoikis, ROCK inhibitor (Y- 27632, Tocris) at a concentration of 10 μM was added to the culture medium for 2 days.

### Radiation response assays

To model radiation resistance, we adapted a previously published medium-throughput radiation dose response assay for compatibility with 3D organoid technology (14,15). In summary, 500 three-day old organoids were embedded in a mixture of BME matrix and culture medium and plated in a 96-well plate as 4 μl droplets, thus allowing growth in 3D structures. Three hours later, organoids were irradiated with a single dose of 0 – 8 Gy using a linear accelerator (Elekta Precise Linear Accelerator 11F49 or Elekta Synergy Agility Linear Accelerator) with a separate plate used for each dose. To allow for photon scattering, the plates were placed on top of a 2-cm polystyrene board. After seven days, cell viability was assessed using CellTiter-Glo 3D (Promega), which is specifically validated for 3D microtissue cultures. The dose response data was normalized to 0 Gy, and then the area under the curve was calculated based on this normalized data. To standardize the area under the curve, it was divided by the maximum possible area that a curve could occupy.

### Bulk genome sequencing

Whole exome sequencing data of HUB015 was obtained from the HUB foundation. Targeted next generation sequencing was performed as described (35). Briefly, 20 ng of DNA extracted from organoid samples was used for sequencing, utilizing the Cancer Hotspot Panel v2 (Thermo Fisher); with some additional hotspots.

Sequencing was carried out on the Ion Chef System (Thermo Fisher) and the IonTorrent S5 sequencer (Thermo Fisher), resulting in an average sequencing depth of 500x.

### Subclonal evolution modelling

To assess subclonal evolution in response to radiation therapy, 250,000 single cells were plated in a 6-well plate and exposed to either 0 or 10 Gy of radiation three days later. In the treatment group (10 Gy), ‘recurrence’ was defined as the regrowth of cancer cells to a sufficient extent that allowed for subsequent passaging of the cells. Organoids were processed for single-cell whole genome sequencing 7 days after the first passage. The control group (0 Gy) was passaged every week.

### Single-cell whole genome sequencing

Organoids were dissociated into single cells with TrypLE (Gibco). After washout of TrypLE, cells were frozen in 500 μl of recovery cell culture freezing medium (Gibco) for subsequent sorting and sequencing. G1 single nuclei, identified by propidium iodide and Hoechst staining, were sorted into a 384-well plate with 10 μl of mineral oil (Sigma-Aldrich) in each well, and stored at −80 °C. Cell lysis was carried out for 2 hours at 50 °C using Proteinase K (Ambion) in 1x Cutsmart (New England Biolabs), followed by heat inactivation at 80 °C for 15 minutes. Genomic DNA was then fragmented with 100 nl of 1 U NlaIII (New England Biolabs) in 1× Cutsmart (New England Biolabs) for 2 hours at 37 °C, followed by heat inactivation at 65 °C for 20 min. Next, the following was added to each well: (i) 50 nl containing 50 mM barcoded double-stranded NLAIII adapters; (ii) 150 nl of 1× T4 DNA ligase buffer containing 40 U T4 DNA ligase (New England Biolabs), supplemented with 10 mM ATP (Invitrogen). The mixture was ligated overnight at 16 °C, after which samples were pooled. Library preparation was performed as previously described (36).

Libraries were sequenced on an Illumina Nextseq 2000 with 2 × 100/150-bp paired-end sequencing.

### Single-cell sequencing data processing

#### Preprocessing and quality selection

The sequencing data was processed using the *snakemake* workflows in Python (v. 3.6), as described previously (37). Briefly, the cell barcodes were extracted and trimmed, and reads without NlaIII sequence or PCR-duplicated reads were removed. The trimmed reads were then mapped to hg38 using BWA 0.7.16a-r1181. R package ‘AneuFinder’ (v. 1.26.0) was used for GC correction, blacklisting of artefact-prone regions and copy number calling (38). Copy numbers were called using the edivisive algorithm with variable width bins (mean 0.5 Mb) based on mappability. Following the removal of cells with high bin-to-bin variation (> 0.7 spikiness) and cells with too few reads, a total of 2,994 high-quality cells remained, exhibiting an average of 26.9 ± 11.7 unique copy number alterations per cell.

#### Clustering of subclones

To create the clustering heat maps, Euclidean distances were calculated from the copy number state measurements with R’s inbuilt ‘dist and ‘hclust’ functions using wrapper functions from package ‘AneuFinder’. For each individual organoid, the matrices containing copy number values before and after radiation were used to perform principal component analysis. Principal component 1 and principal component 2 were plotted on the x and y axes, respectively. As such, each dot in the principal component plots represents a single-cell copy number profile and is colored either by radiation status or by subclone group. The optimum number of subclones was defined by performing k-means clustering on the first 10 principal components and selecting the maximum average silhouette (*s*) width:

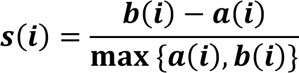

Where *a(i)* is the average intra-cluster distance and *b(i)* is the average inter-cluster distance. Clusters that were separated on a principal component explaining less than 10% of the variance were dismissed as true subclones (e.g., HUB183), or labeled as minor subclones if they were clearly separated (e.g., HUB015). Using this approach, the following k values were used: HUB183 (k = 1), HUB005 (k = 3), HUB015 (k = 3), HUB106 (k = 3), HUB062 (k = 3), and HUB197 (k = 3).

Subclonal expansion was defined as a significant 10-fold increase in the relative dominance of subclones, accompanied by the rejection of the null hypothesis of Fisher’s exact test (thus indicating that the relative proportions of the subclones are dependent on the radiation status). Subclonal extinction referred to the total disappearance of subclones. Finally, subclonal persistence was defined as the maintenance of subclonal composition with minimal changes (less than 10-fold) before and after radiation.

#### Resistance-specific copy number analysis

Consensus copy number profiles of subpopulations (either whole organoids or subclones within organoids) were calculated using custom scripts. A copy number alteration was considered as ‘consensus’ when present in more than 60-80% of cells. To identify resistance-specific copy number alterations, consensus profiles of resistant subclones were compared to consensus profiles of sensitive subclones.

Resistant subclones were defined as subclones that either persisted or expaned upon irradiation. To ensure specificity, a copy number alteration was deemed ‘resistance-specific’ only if present in more than 80% of resistant subclones, and in less than 10% of the sensitive subclones. All subclones within HUB015 (diploid) and whole-genome duplicated subclones within HUB197 were excluded from this analysis. COSMIC cancer gene census oncogenes were downloaded from https://cancer.sanger.ac.uk/census and mapped to each copy number alteration (39).

#### Copy number pattern analysis

Heterogeneity was defined as described previously (40):

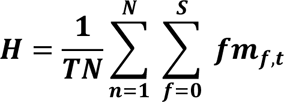

where T and N sets containing bins and single cells, respectively. S represents the total number of possible copy number states, while ***m***_*f,t*_ denotes the number of cells with a particular copy number s at bin t. Copy number transitions were shifted to a single shared middle position, thereby avoiding an overestimation of heterogeneity due to technical issues that cause shifts of copy number states transitions (40).

For copy number length analysis, each copy number alteration was standardized by dividing its length by the length of its corresponding chromosome arm. As such, a value of 1 indicated an alteration spanning an entire arm, while 2 indicated a whole-chromosome alteration. If the alteration spanned the centromere, the fractions for each arm were summed. Alterations were categorized as focal if their length value was less than 0.3, as large if their length value was between 0.3 and 0.98, and as arm to whole-chromosome level if their length value was greater than 0.98 (41).

### Chromosomal instability signature

Segment level copy number data (‘CCLE_segment_cn.csv’) from the Cancer Cell Line Encyclopedia (CCLE) were downloaded from https://depmap.org (27,42). We used the 22Q2 DepMap release. Chromosomal instability signatures scores were computed using the ‘quantifyCNSignatures’ as described previously (26). This wrapper function computes signature scores for 17 chromosomal instability signatures, each having a unique putative cause (26). *In vitro* radiation response data of CCLE cell lines were obtained from Yard et al. (15). To compare signature scores between top radioresistant and radiosensitive cell lines the following linear model was fitted using the ‘lm’ package in R:

[umath3]

### Statistical analysis

R version 4.2.1 was used for statistical analysis. Continuous variables were compared using the Mann–Whitney *U* test or Student’s *t* test, where appropriate. Categorical variables were compared using Fisher’s exact test. In the case of multiple comparisons *P* values were adjusted using the Bonferroni procedure.

### Data Availability

The raw sequencing data from this study are available in the NCBI Sequence Read Archive under accession number PRJNA1015247. Processed data will be shared by the lead contact upon reasonable request.

### Code Availabilityw

Code to reproduce the analyses and figures is available at https://github.com/dshandel/subclonal_evolution_paper

---

**Figure S1.**
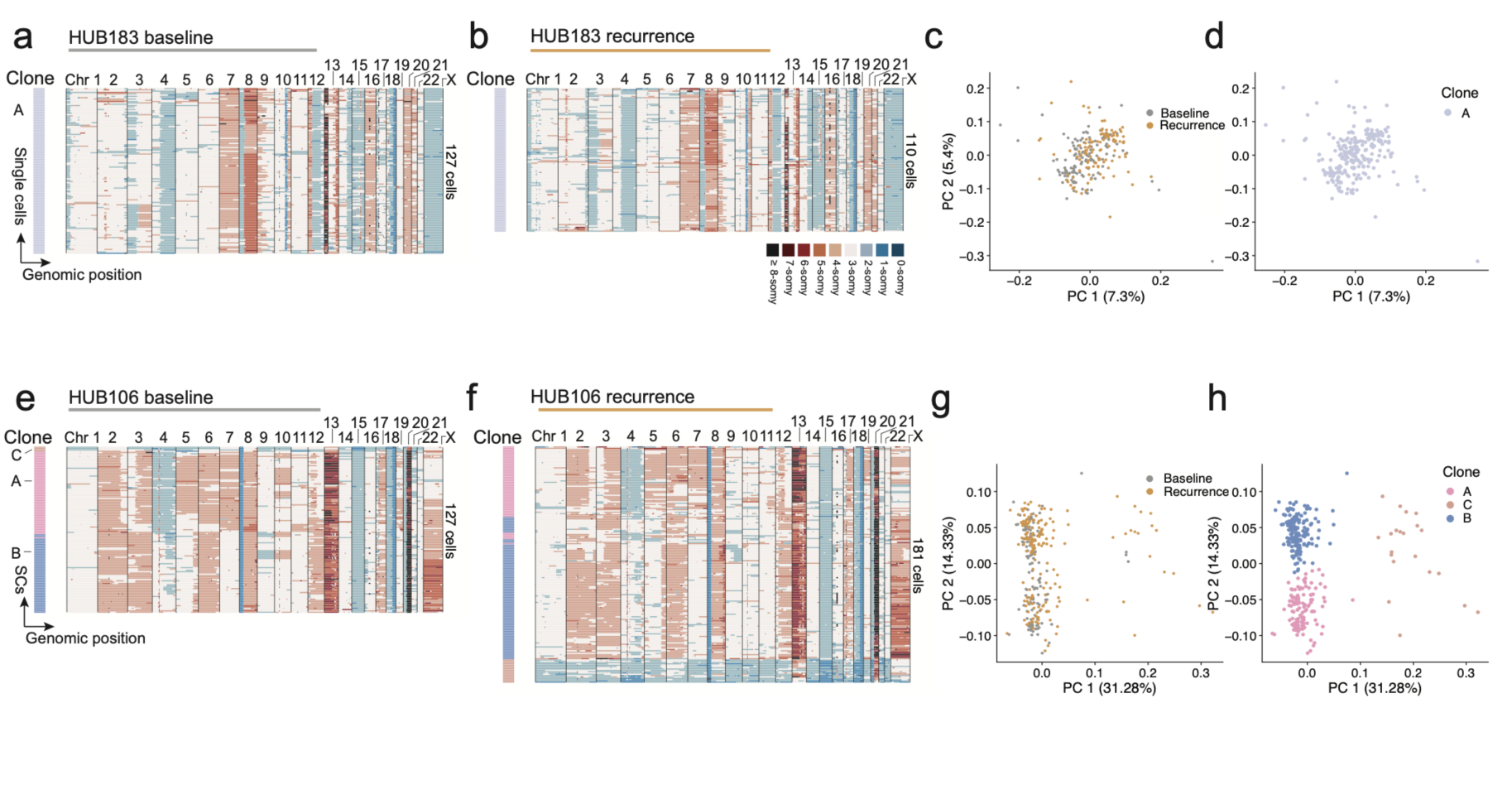
Subclonal evolution in persister organoids. a, b, Single cell copy number heatmaps of HUB183 (resistant, persister) at baseline (**a**) and after recurrence (**b**). **c, d,** PCA plots of single cell copy number profiles from the baseline and recurrence populations. Colors indicate treatment status (**c**) or subclone (**d**) as derived from k- means clustering. **e, f,** Cluster heatmaps of single cell copy number profiles of HUB106 (sensitive, persister) at baseline (**e**) and after recurrence following 10 Gy (**f**). **g, h,** PCA plots of single cell copy number profiles from the baseline and recurrence populations. SCs: single cells.

**Figure S2.**
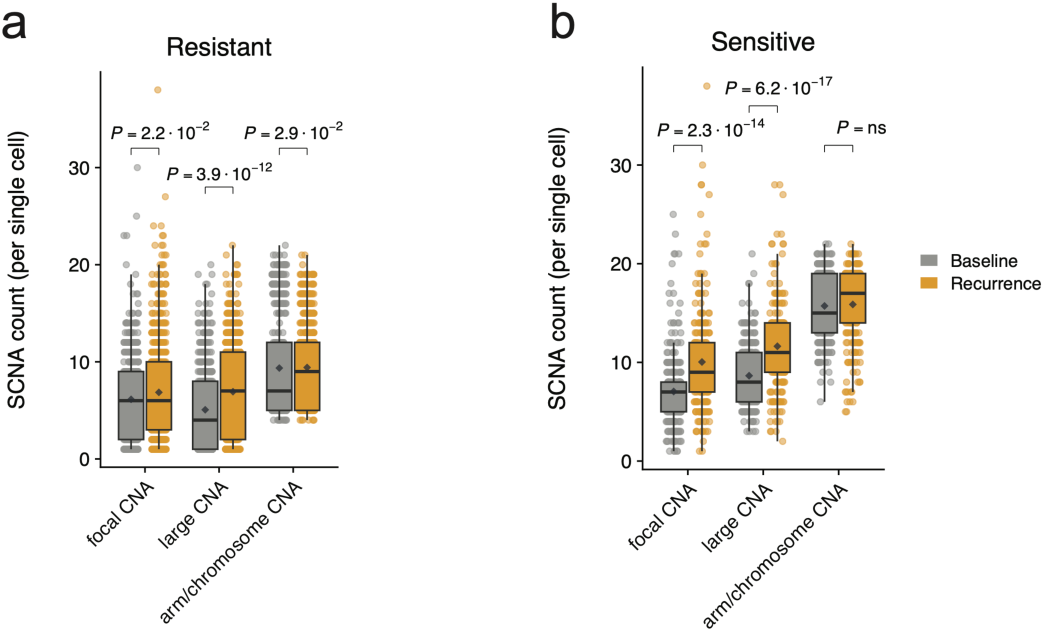
Somatic copy number alterations in resistant and sensitive single cells a, b, Somatic copy number alterations count in single cells from resistant (**a**) and sensitive (**b**) subclones at baseline and recurrence. Copy number alterations are stratified into focal, large and arm to whole-chromosome lengths (S)CNA: (somatic) copy number alterations. *P*-values from Mann-Whitney *U* test.

**Figure S3.**
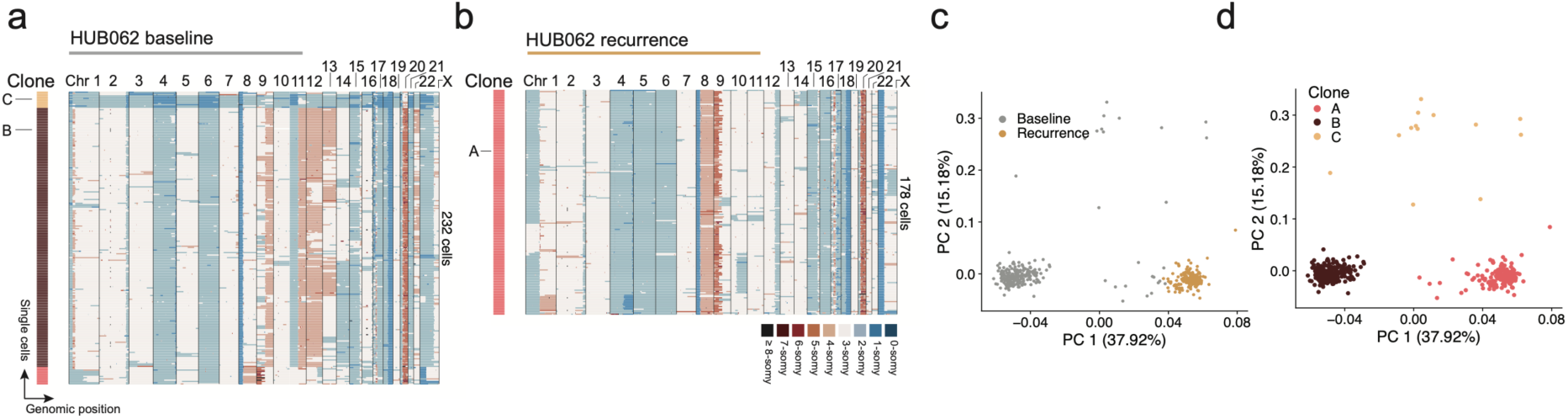
Subclonal evolution in HUB062. a, b, Single cell copy number heatmaps of HUB062 at baseline (**a**) and after recurrence (**b**). **c, d,** PCA plots of single cell copy number profiles from the baseline and recurrence populations. Colors indicate treatment status (**c**) or subclone (**d**).

